# Sulfur disproportionating microbial communities in a dynamic, microoxic-sulfidic karst system

**DOI:** 10.1101/2023.03.26.534238

**Authors:** Heidi S. Aronson, Christian E. Clark, Douglas E. LaRowe, Jan P. Amend, L. Polerecky, Jennifer L. Macalady

**Author notes:** Corresponding author: Jennifer Macalady 210 Deike Building University Park, PA, USA, 16802. These authors contributed equally to the study (Heidi S. Aronson, Christian E. Clark, Jennifer L. Macalady). **Competing interests:** The authors declare no competing interests.

## Abstract

Biogeochemical sulfur cycling in sulfidic karst systems is largely driven by abiotic and biological sulfide oxidation, but the fate of elemental sulfur (S^0^) that accumulates in these systems is not well understood. The Frasassi Cave system (Italy) is intersected by a sulfidic aquifer that mixes with small quantities of oxygen-rich meteoric water, creating Proterozoic-like conditions and supporting a prolific ecosystem driven by sulfur-based chemolithoautotrophy. To better understand the cycling of S^0^ in this environment, we examined the geochemistry and microbiology of sediments underlying widespread sulfide-oxidizing mats dominated by *Beggiatoa*. Sediment populations were dominated by uncultivated relatives of sulfur cycling chemolithoautotrophs related to *Sulfurovum*, *Halothiobacillus, Thiofaba, Thiovirga, Thiobacillus*, and *Desulfocapsa*, as well as diverse uncultivated anaerobic heterotrophs affiliated with *Bacteroidota*, Anaerolineaceae, Lentimicrobiaceae, and Prolixibacteraceae. *Desulfocapsa* and *Sulfurovum* populations accounted for 12-26% of sediment 16S rRNA amplicon sequences and were closely related to isolates which carry out autotrophic S^0^ disproportionation in pure culture. Gibbs energy (Δ*Gr*) calculations revealed that S^0^ disproportionation under *in situ* conditions is energy yielding. Microsensor profiles through the mat-sediment interface showed that *Beggiatoa* mats consume dissolved sulfide and oxygen, but a net increase in acidity was only observed in the sediments below. Together, these findings suggest that disproportionation is an important sink for S^0^ generated by microbial sulfide oxidation in this oxygen-limited system and may contribute to the weathering of carbonate rocks and sediments in sulfur-rich environments.

## Introduction

Actively forming sulfidic caves are valuable natural laboratories for studying microbial ecosystems that are supported entirely by *in situ* chemolithotrophic sulfur cycling and primary productivity. These restricted environments provide constrained geochemical contexts in which to investigate the role of microorganisms in mediating the geochemistry of subsurface environments and their contributions to sulfur cycling and cave formation. The karst at Frasassi (Italy) is a rare example of a cave system actively forming by sulfuric acid speleogenesis (SAS; Table 1, Rxn 1) (Galdenzi *et al*., 1999; Macalady *et al*., 2019). The Frasassi cave system is intersected by a perennially sulfidic (up to 600 µM total dissolved sulfide) deep aquifer. Sulfide that degasses from the water table reacts with oxygen that is either in the downward percolating meteoric water or in the cave air, to form sulfuric acid, which corrodes subaerial carbonate host rock to form gypsum and carbonic acid (Rxn 1).

**Table 1:** Selected chemical and catabolic reactions and their average Gibbs energy yields, ΔGr, under in situ conditions along the microsensor depth profile through Frasassi Beggiatoa mats and sediments shown in Fig. 1. Chemical concentration data that were used to calculate activities can be found in Supplementary File 1. DNRA = dissimilatory nitrate reduction to ammonium.

| Reaction number | Reaction name | Reaction | $\Delta G_r$<br>(kJ/mol S consumed) | $\Delta G_r$<br>(kJ/mol O<sub>2</sub>) | $\Delta G_r$<br>(kJ/mol H<sub>2</sub>) |
| --- | --- | --- | --- | --- | --- |
| 1 | Sulfuric acid speleogenesis | $\text{H}_2\text{SO}_4 + \text{CaCO}_3 + 2\text{H}_2\text{O} \rightleftharpoons \text{CaSO}_4 \cdot 2\text{H}_2\text{O} + \text{H}_2\text{CO}_3$ | | | |
| 2 | Incomplete aerobic sulfide oxidation | $2\text{HS}^- + \text{O}_2 + 2\text{H}^+ \rightleftharpoons 2\text{S}^0 + 2\text{H}_2\text{O}$ | -283.1 | -411.2 | |
| 3 | Complete aerobic sulfide oxidation | $\text{HS}^- + 2\text{O}_2 \rightleftharpoons \text{SO}_4^{2-} + \text{H}^+$ | -761.0 | -381.6 | |
| 4 | Aerobic S<sup>0</sup> oxidation | $2\text{S}^0 + 3\text{O}_2 + 2\text{H}_2\text{O} \rightleftharpoons 2\text{SO}_4^{2-} + 4\text{H}^+$ | -583.3 | -383.3 | |
| 5 | Incomplete sulfide oxidation coupled to DNRA | $4\text{HS}^- + \text{NO}_3^- + 6\text{H}^+ \rightleftharpoons 4\text{S}^0 + \text{NH}_4^+ + 3\text{H}_2\text{O}$ | -89.0 | | |
| 6 | Complete sulfide oxidation coupled to DNRA | $\text{HS}^- + \text{NO}_3^- + \text{H}_2\text{O} + \text{H}^+ \rightleftharpoons \text{SO}_4^{2-} + \text{NH}_4^+$ | -606.2 | | |
| 7 | S<sup>0</sup> oxidation coupled to DNRA | $4\text{S}^0 + 3\text{NO}_3^- + 7\text{H}_2\text{O} \rightleftharpoons 4\text{SO}_4^{2-} + 3\text{NH}_4^+ + 2\text{H}^+$ | -331.8 | | |
| 8 | Incomplete sulfide oxidation coupled to denitrification | $5\text{HS}^- + 2\text{NO}_3^- + 7\text{H}^+ \rightleftharpoons 5\text{S}^0 + \text{N}_2 + 6\text{H}_2\text{O}$ | -174.0 | | |
| 9 | Complete sulfide oxidation coupled to denitrification | $5\text{HS}^- + 8\text{NO}_3^- + 3\text{H}^+ \rightleftharpoons 5\text{SO}_4^{2-} + 4\text{N}_2 + 4\text{H}_2\text{O}$ | -735.5 | | |
| 10 | S<sup>0</sup> oxidation coupled to denitrification | $5\text{S}^0 + 6\text{NO}_3^- + 2\text{H}_2\text{O} \rightleftharpoons 5\text{SO}_4^{2-} + 3\text{N}_2 + 4\text{H}^+$ | -583.2 | | |
| 11 | S<sup>0</sup> disproportionation | $4\text{S}^0 + 4\text{H}_2\text{O} \rightleftharpoons 3\text{HS}^- + \text{SO}_4^{2-} + 5\text{H}^+$ | -16.2 | | |
| 12 | S<sup>0</sup> reduction | $\text{S}^0 + \text{H}_2 \rightleftharpoons \text{HS}^- + \text{H}^+$ | -25.6 | | -25.6 |
| 13 | Sulfate reduction | $\text{SO}_4^{2-} + 4\text{H}_2 + \text{H}^+ \rightleftharpoons \text{HS}^- + 4\text{H}_2\text{O}$ | -37.3 | | -9.3 |
| 14 | S<sub>2</sub><sup>2-</sup> disproportionation | $4\text{S}_2^{2-} + 4\text{H}_2\text{O} \rightleftharpoons 7\text{HS}^- + \text{SO}_4^{2-} + \text{H}^+$ | -49.9 | | |
| 15 | S<sub>3</sub><sup>2-</sup> disproportionation | $2\text{S}_3^{2-} + 4\text{H}_2\text{O} \rightleftharpoons 5\text{HS}^- + \text{SO}_4^{2-} + 3\text{H}^+$ | -18.4 | | |
| 16 | S<sub>4</sub><sup>2-</sup> disproportionation | $4\text{S}_4^{2-} + 12\text{H}_2\text{O} \rightleftharpoons 13\text{HS}^- + 3\text{SO}_4^{2-} + 11\text{H}^+$ | -16.7 | | |
| 17 | Ferric iron reduction | $\text{HS}^- + 2\text{FeOOH} + 5\text{H}^+ \rightleftharpoons 2\text{Fe}^{2+} + \text{S}^0 + 4\text{H}_2\text{O}$ | | | |
| 18 | Iron sulfide formation | $\text{HS}^- + 2\text{Fe}^{2+} \rightleftharpoons \text{FeS} + 4\text{H}^+$ | | | |
| 19 | Sulfur disproportionation with ferric iron as a sulfide scavenger | $3\text{S}^0 + 2\text{FeOOH} \rightleftharpoons 2\text{FeS} + \text{SO}_4^{2-} + 2\text{H}^+$ | | | |

In the water table, the mixing of oxygenated meteoric water with the sulfidic aquifer forms anoxic, microoxic, and oxygenated zones (Galdenzi *et al*., 2008; Macalady *et al*., 2008; Meyer & Kump, 2008; Lyons *et al*., 2009). White microbial mats form at the interface between the microoxic stream waters and underlying sediments. These mats are dominated by chemolithotrophic sulfur-cycling taxa, including freshwater *Beggiatoa* strains, Campylobacterota (f.k.a. Epsilonproteobacteria), Burkholderiales (f.k.a. Betaproteobacteria), and Desulfobacterota. The Frasassi stream microbial mats have been well described (Macalady *et al*., 2006, 2008; Hamilton *et al*., 2015; Jones *et al*., 2015). The subaqueous *Beggiatoa* mats primarily oxidize sulfide, and abundant visible sulfur inclusions within their cells suggest that sulfide is oxidized incompletely to form elemental sulfur (S^0^, Table 1, Rxn 2) (Macalady *et al*., 2006, 2008; Hamilton *et al*., 2015). Sulfur speciation measurements on Frasassi microbial mats using K-edge X-ray absorption near-edge structure spectroscopy confirmed that the mats contained S^0^, primarily in the form of S8 (Engel *et al*., 2007). Additionally, microsensor measurements through the mats did not detect a decrease in pH, indicating that incomplete oxidation of sulfide, rather than complete oxidation to sulfuric acid, was occurring (Table 1, Rxn 3; Fig. 1; Jones *et al*., 2015).

**Figure 1:**
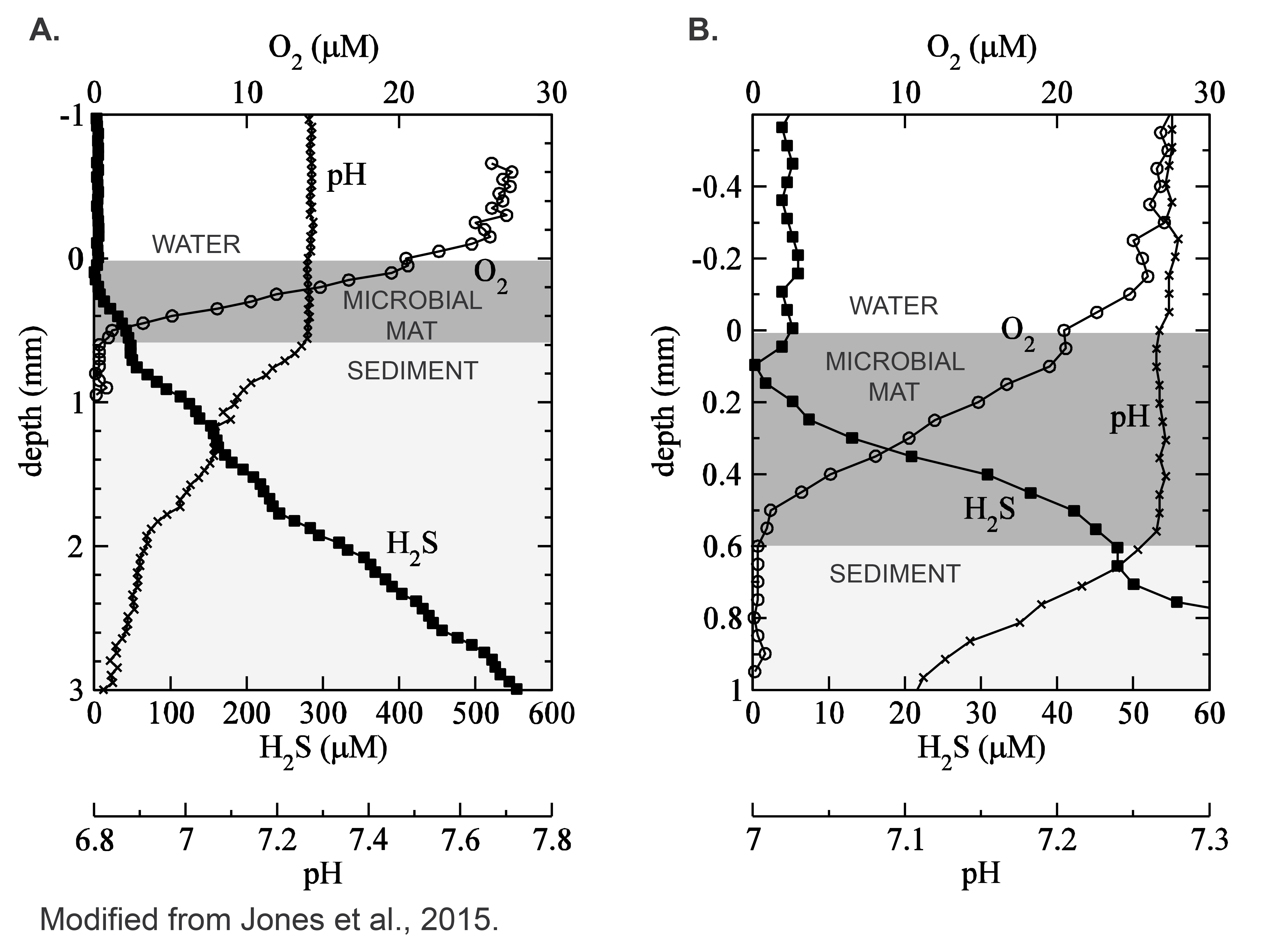
Representative *in situ* concentration profiles of H2S, O2, and pH as a function of depth through a Frasassi Cave *Beggiatoa* mat and sediment layer at Pozzo dei Cristalli. Panel B is a magnified version of panel A.

In contrast to the mats, net acid and sulfide production was detected in the sediments, suggesting that S^0^ is being used as an electron donor and/or acceptor (Fig 1., Jones *et al*., 2015). There are four potential fates of S^0^ in the Frasassi sediments, including 1) physical transport (export from the cave system by flowing water), 2) microbial oxidation, 3) microbial reduction, and 4) microbial disproportionation. There is currently no good estimate of S^0^ export, but S^0^ can be observed exiting the cave system into the Sentino River. Here, we explored possible microbial roles in S^0^ consumption by analyzing the microbial communities within the sediments and calculating the energetic yield of reactions involving S^0^ using *in situ* water chemistry.

## Materials and Methods

### Site description, geochemistry, and sample collection

The stream waters at Frasassi are circumneutral (pH 6.9-7.4) and have a nearly constant temperature of 13-14°C, total sulfide concentrations up to 600 µM, and conductivity between 1000-3500 µS/cm, with an ionic strength equivalent to a salinity of approximately 3.5-4.5 ppt (Galdenzi *et al*., 2008; Macalady *et al*., 2008). The stream waters have consistently low levels of dissolved oxygen (<30 µM), nitrate concentrations below detection limits (∼7 nM), and ammonium concentrations between 30 and 175 µM (Macalady *et al*., 2006, 2008).

*Beggiatoa* mat and sediment samples for geochemical and microbial community analysis were collected in 2006, 2007, and 2010 from sulfidic streams in the Frasassi cave system, specifically from Ramo Sulfureo (RS), Grotta Sulfurea (GS), Pozzo dei Cristalli (PC), and Vecchio Condotto (VC) (Table 2; for details on sample locations see (Macalady *et al*., 2006, 2007, 2008; Hamilton *et al*., 2015)). Paired mat- sediment samples were collected by using a sterile transfer pipette to remove the white mat into a sterile 50 mL centrifuge tube. The underlying sediment was then collected with a new transfer pipette and placed into a sterile 50 mL centrifuge tube. All samples were preserved by adding 3:1 RNAlater:sample volume (Sigma-Aldrich Corp., St. Louis, MO), and stored at -20°C until processing at Pennsylvania State University in 2015.

**Table 2:**
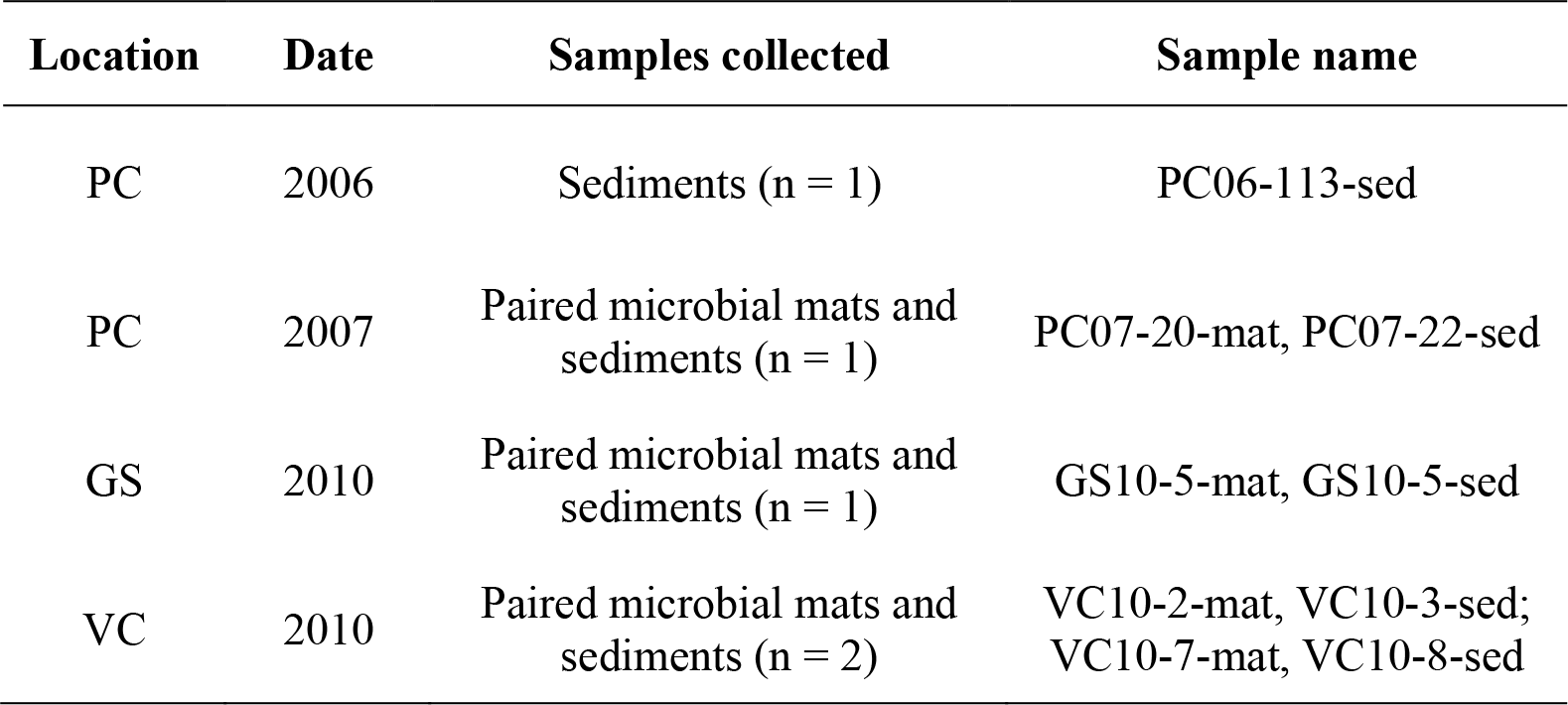
Samples collected for 16S rRNA gene sequencing. Location names refer to Pozzo dei Cristalli (PC), Grotta Sulfurea (GS), and Vecchio Condotto (VC).

### Reaction energetics

The Gibbs energy yields (Δ*Gr*) for various sulfur redox reactions (Table 1) were calculated with

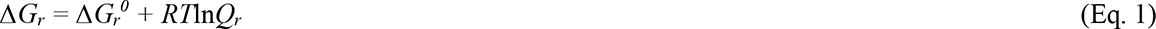

where Δ*Gr*^0^ is the standard state Gibbs energy, *R* is the universal gas constant, T is the temperature in Kelvin, and *Qr* refers to the reaction activity quotient. Values of Δ*Gr*^0^ were calculated at 13.4 °C and 1 bar with the revised Helgeson-Kirkham-Flowers (HKF) equations of state (Helgeson *et al*., 1981; Tanger & Helgeson, 1988; Shock *et al*., 1992) using the “subcrt” command from the R software package CHNOSZ v1.4.1 (Dick, 2019). Thermodynamic data in CHNOSZ are derived from the OrganoBioGeoTherm database, which come from a number of sources (https://chnosz.net/download/refs.html). The sources of these data are provided in the CHNOSZ package documentation. Values of *Qr* were calculated with

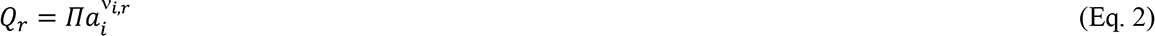

where *ai* represents the activity of the *i*^th^ species raised to its stochiometric reaction coefficient *νi,r*, in the *r*^th^ reaction, which is positive for products and negative for reactants.

Activities were calculated with the relation

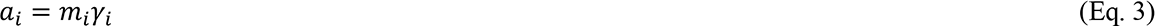

where *mi* and *γi* are the molality and activity coefficients of the *i*^th^ species. Concentrations were sourced from median values from Pozzo dei Cristalli stream water data collected from 2002-2011 and were converted to molality by assuming stream water density of 1 kg/L. (Supplemental File 1). Concentrations of sulfide, oxygen, and protons were sourced from microsensor data from Jones *et al*. 2015 (Fig. 1, Supplemental File 1). Oxygen concentrations below the detection limit (≤2 µM) for depths below 0.5 mm were set between 10^-6^ molal and 10^-8^ molal to illustrate the effects of extremely low oxygen concentration on Gibbs energy yields (Supplemental File 1). H_2_ concentrations were assumed to range between 10^-10^ to 10^-8^ molal (Hoehler *et al*., 1998) and polysulfide concentrations were assumed to be 10^-5^ molal. Activities for aqueous species were determined using the aqueous speciation package AqEquil v0.9.1 (Boyer *et al*., 2021), which is based on EQ3/6 (Wolery, 1979). The activities of elemental sulfur and water were assumed to be unity (*ai* = 1). Values of Δ*Gr* were calculated for various redox reactions involving sulfur in various oxidation states (see Table 1) along the depth profile shown in Fig. 1. The Gibbs energy for S^0^ disproportionation was calculated for sulfate and sulfide concentrations ranging from 1x10^-9^ molal to 3x10^- 3^ or 1x10^-3^ molal, respectively, with pH set at 7.26 and a temperature of 13.4 °C (median pH and temperature of Frasassi waters).

### 16S rRNA gene amplicon sequencing and analysis

Replicate DNA extractions from samples and blanks were performed in 2015 in a clean fume hood using MoBio Power Lyzer Soil DNA isolation kit #12855-50 according to kit protocol (Mo Bio Laboratories Inc., Carlsbad, CA). The microbial community of each sample was analyzed by 16S rRNA gene amplicon sequencing of the V4-V5 hypervariable region (Illumina MiSeq platform) (Illumina Inc., San Diego, CA) using 515F and 806R universal primers at the MrDNA Laboratory (Molecular Research LP, Shallowater, TX). The primers were used in a 28 cycle PCR with the HotStarTaq Plus Master Mix Kit (Qiagen Inc., Hilden, Germany). The PCR cycle conditions were as follows: 94 °C for 3 minutes, succeeded by 28 cycles at 94 °C for 30 seconds, 53 °C for 40 seconds and 72 °C for 1 minute. A final elongation step was carried out at 72 °C for 5 minutes. PCR products were assessed in 2% agarose gel to determine amplification success. Pooled samples were purified using calibrated Ampure XP beads. Purified PCR products were then used to compile the Illumina DNA library. Sequence data were processed using the MrDNA pipeline (Molecular Research LP, Shallowater, TX). Forward and reverse reads were joined, their barcodes were removed, sequences with less than 150 base pairs were removed, and ambiguous base calls were removed.

16S rRNA amplicon sequence data were processed using QIIME2 v2019.7.0 (Bolyen *et al*., 2019). Fastq data were imported into QIIME2 using the fastq manifest format. Sequences were demultiplexed and denoised into amplicon sequence variants (ASVs) using the QIIME2 DADA2 “denoised-paired” plugin (Callahan *et al*., 2016). Taxonomy of each ASV was assigned using the “feature-classfier classify-sklearn” plugin. The classifier was trained on the SILVA138 database clustered at 99% and trimmed to the amplified region (Quast *et al*., 2012). Mitochondrial sequences were removed using “taxa filter-table” plugin. The R package phyloseq was used to further analyze and visualize data from the filtered OTU table and taxonomy (McMurdie & Holmes, 2013). Sequences that were unassigned at the domain level were removed and sequence counts were transformed to relative abundance. Plots were produced in R using the package ggplot2 (Wickham, 2016; R Core Team, 2019). Interpretations of microbial metabolic potential were inferred from metabolisms of the closest cultured relatives within the same genus and from previous metagenomic studies of Frasassi biofilms (Tsao, 2014; Hamilton *et al*., 2015; McCauley Rench, 2015; Labrado, 2017; Pavia *et al*., 2018).

Phylogenetic trees were constructed using *Desulfocapsa, Sulfurovum,* and Halothiobacillaceae 16S rRNA gene sequences from this study and from close relatives. *Desulfocapsa, Sulfurovum,* and Halothiobacillaceae sequences from this study were blasted against all 16S rRNA sequences from previous Frasassi studies (Macalady *et al*., 2006, 2007, 2008; Jones *et al*., 2008, 2010, 2012, 2014, 2016; Hamilton *et al*., 2015), against the NCBI 16S rRNA gene sequence reference database, and against the NCBI nt sequence database. The Halothiobacillaceae tree also contains additional *Thiobacillus* sequences from this study. Frasassi sequences ≥97% similar to sequences from this study, reference sequences ≥90% similar to sequences from this study, and nt sequences 100% similar to sequences from this study were included in the trees. The trees were pruned by selecting representative sequences with the dist.seqs, cluster (furthest neighbor at most 2% distant), and get.oturep commands in mothur (Schloss *et al*., 2009). Sequences were aligned using MUSCLE (Edgar, 2004) and trimmed with trimal (Capella-Gutiérrez *et al*., 2009). Phylogenetic trees were constructed using IQ-TREE with a maximum-likelihood method and the generalized time-reversible model of nucleotide evolution with 1000 bootstrap replications (Nguyen *et al*., 2015).

## Results

### Reaction energetics

Gibbs energies of the reactions listed in Table 1 are shown in Table 1, Fig. 2, and Fig. S4. For S^0^ disproportionation and reduction, sulfate reduction, and polysulfide disproportionation (Table 1, Rxns. 11- 16), values of Δ*Gr* along the microsensor depth profile were negative (exergonic) (Fig. 2). The Gibbs energy yields for these reactions became less negative with greater depth because of increasing sulfide concentration and reached local negative maxima at 0.1 mm depth because of the minimum sulfide concentration (1 µM). Values of Δ*Gr* along the depth profile different substantially among the different sulfur species that were disproportionated or reduced, with S2^2-^ disproportionation yielding the most energy at up to -55 kJ mol^-1^ S and S^0^ disproportionation yielding the least energy at as low as -13 kJ mol^-1^ S (Fig. 2). Values of Δ*Gr* for S^0^ disproportionation were between -8 and -25 kJ mol^-1^ S for the range of sulfate and sulfide activities found at Frasassi (Fig. S4). The oxidation reactions with oxygen or nitrate yield significantly more energy (-91 to -763 kJ mol^-1^ S) even with activities equivalent to oxygen concentrations 10^-8^ molal (Table 1, Rxns. 1-10, Supplemental File 1).

**Figure 2:**
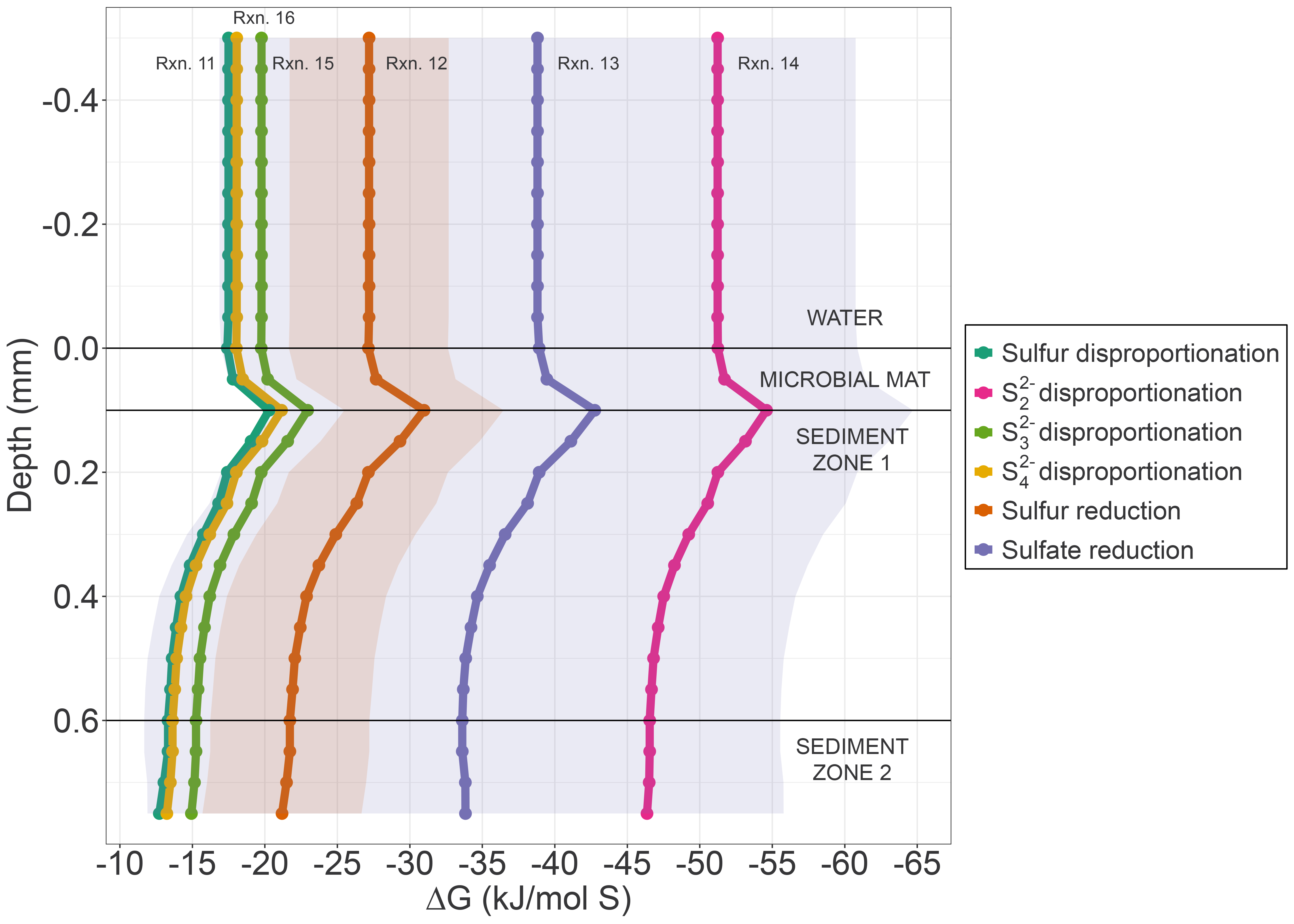
Gibbs energies, Δ*Gr*, of S^0^ disproportionation, S^0^ reduction, sulfate reduction, and polysulfide disproportionation (reactions 11-16 in Table 1) as a function of depth into Frasassi cave sediment at T = 13.5°C and pH = 7.26 expressed in units of kJ/mol S. Activities of sulfide, oxygen, and protons were derived from microsensor data shown in Fig. 1B. Dots correspond to Δ*Gr* at each point on the microsensor data from Fig. 1B, solid lines designate guiding fit lines, and the colored ribbons around these lines indicate the energetic yield when hydrogen concentrations were set at one order of magnitude higher and lower than the values used for the plotted lines (1 nM) for S^0^ reduction (orange) and sulfate reduction (purple). Reaction numbers on the plot refer to reactions listed in Table 1.

### Microbial community composition

Analysis of the 16S rRNA gene V4-V5 hypervariable region revealed similar community compositions for each sample type across locations and sampling times (Fig. 3). Taxonomy of each ASV was identified at the genus level, and interpretations of potential metabolic functions were inferred from the metabolisms of the closest cultured relatives and from previous metagenomic studies (Tsao, 2014; Hamilton *et al*., 2015; McCauley Rench, 2015; Labrado, 2017; Pavia *et al*., 2018). Sequences were affiliated primarily with Bacteria; Archaea represented fewer than 3% of reads from any sample and typically fewer than 1%.

**Figure 3:**
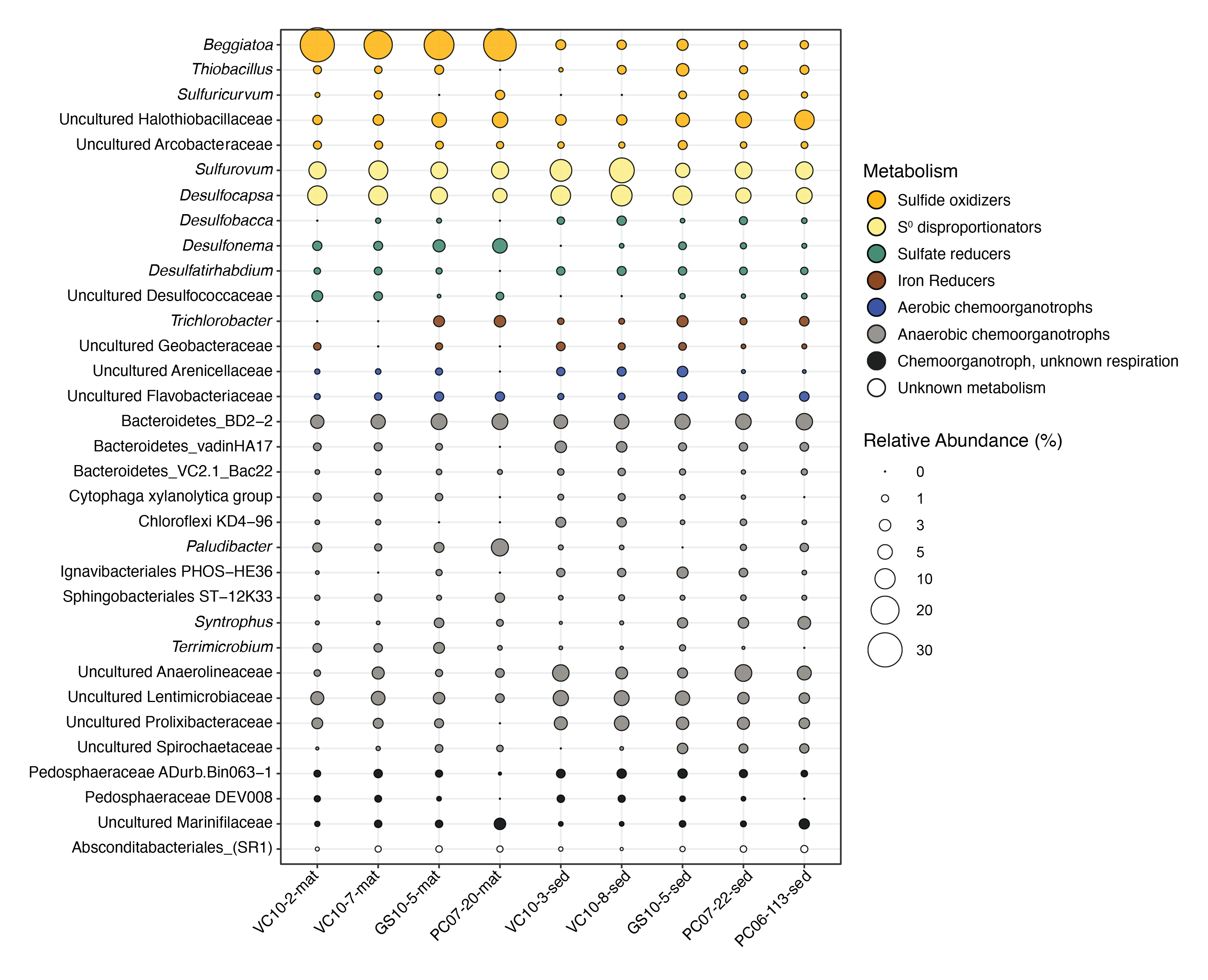
Abundance and inferred catabolisms of microbial taxa from paired Frasassi biofilm and sediment samples. Catabolisms were assigned based on published accounts of closely related species and data from the current study. Only taxonomic affiliations of Illumina V4 amplicons that comprised >0.5% of genera were included. Some strains related to the sulfide oxidizers and disproportionators shown can also reduce S^0^ using H2, e.g. *Sulfurovum, Sulfuricurvum,* and *Geobacter*.

All mat samples were dominated by the sulfur-oxidizing Gammaproteobacterial genus *Beggiatoa* (20-30%), consistent with field and microscopic observations of gliding filaments coiled at the sediment surface. *Beggiatoa* sequences were significantly lower in relative sequence abundance (<3%) in anoxic sediment immediately below the mat surface. The remaining populations in all samples were dominated by *Sulfurovum,* (Campylobacterota, f.k.a. Epsilonproteobacteria, 5-15%), uncultured strains of Halothiobacillaceae (Gammaproteobacteria, 2-9%)*, Desulfocapsa* (Desulfobacterota, 5-11%), and Bacteroidetes BD2-2 (4-7%). Other anaerobic chemoorganotrophs, including Bacteroidetes vadinHA17, uncultured Anaerolineaceae, uncultured Lentimicrobiaceae, uncultured Prolixibacteraceae, and Pedosphaeraceae ADurb.Bin063-1 were consistently higher in abundance in the sediments than in the mats. There were 6 clades of Frasassi *Sulfurovum* sequences that were most closely related to sulfide-oxidizing taxa *Nitratifractor salsuginis*, *Sulfurovum aggregans, Sulfurovum lithotrophicum,* and *Sulfurovum riftiae,* with the most abundant sequences in clade 4 (Fig. S5). Reads affiliated with uncultured Halothiobacillaceae made up 2-10% of the sediment and mat communities (Fig. 3). There were three clades of sequences associated with the Halothiobacillaceae (Fig. S6). One clade clustered with *Halothiobacillus neapolitanus*, one clustered with *Thiovirga* sequences, and the third, which included the most abundant sequences, clustered with *Thiofaba tepidiphila*. There were two clades of *Thiobacillus* from Frasassi sequences, which both clustered with *Thiobacillus thioparus, Thiobacillus sanjanensis, Thiobacillus denitrificans, and Thiobacillus thiophilus* (Fig. S6). *Desulfocapsa* sequences made up 5-11% of total reads (Fig 3). The majority (n =19) of *Desulfocapsa* strains from this study were most closely related to *Desulfocapsa thiozymogenes* (Fig. S7). Other abundant (>1%) populations were associated with other sulfur-oxidizing taxa (*Thiobacillus, Sulfuricurvum,* and *Arcobacteraceae*), sulfate-reducing taxa (*Desulfobacca, Desulfonema,* and *Desulfatirhabdium*), and anaerobic chemoorganotrophs (uncultured Anaerolinaceae, Lentimicrobiaceae, and Prolixibacteraceae) (Fig. 3).

## Discussion

### S^0^ is produced by dominant members of the microbial mat community

The microoxic, sulfidic streams in the Frasassi karst host perennially abundant white sulfide-oxidizing microbial mats that develop at the water-sediment interface (Macalady *et al*., 2006, 2008). The schematic shown in Fig. 4 provides an overview of our current understanding of the biogeochemistry and microbial ecology governing sulfur cycling in the Frasassi mats and sediments. The *Beggiatoa*-dominated mats consume sulfide and oxygen without a concomitant decrease in pH (Fig. 1, Jones *et al*., 2015), which is consistent with incomplete microbial sulfide oxidation to S^0^ within the mat (Table 1, Rxn 2). A decrease in pH would be expected if complete sulfide oxidation (Table 1, Rxn 3) was occurring (Jørgensen & Revsbech, 1983; Schwedt *et al*., 2012). Dominant members of the mat community, including *Beggiatoa, Sulfurovum,* and Halothiobacillaceae show the genetic potential to perform incomplete and complete sulfide oxidation, suggesting that the end product of sulfide oxidation may be determined by both environmental and physiological factors as discussed in the following (Tsao, 2014; Hamilton *et al*., 2015; McCauley Rench, 2015; Pavia *et al*., 2018).

**Figure 4:**
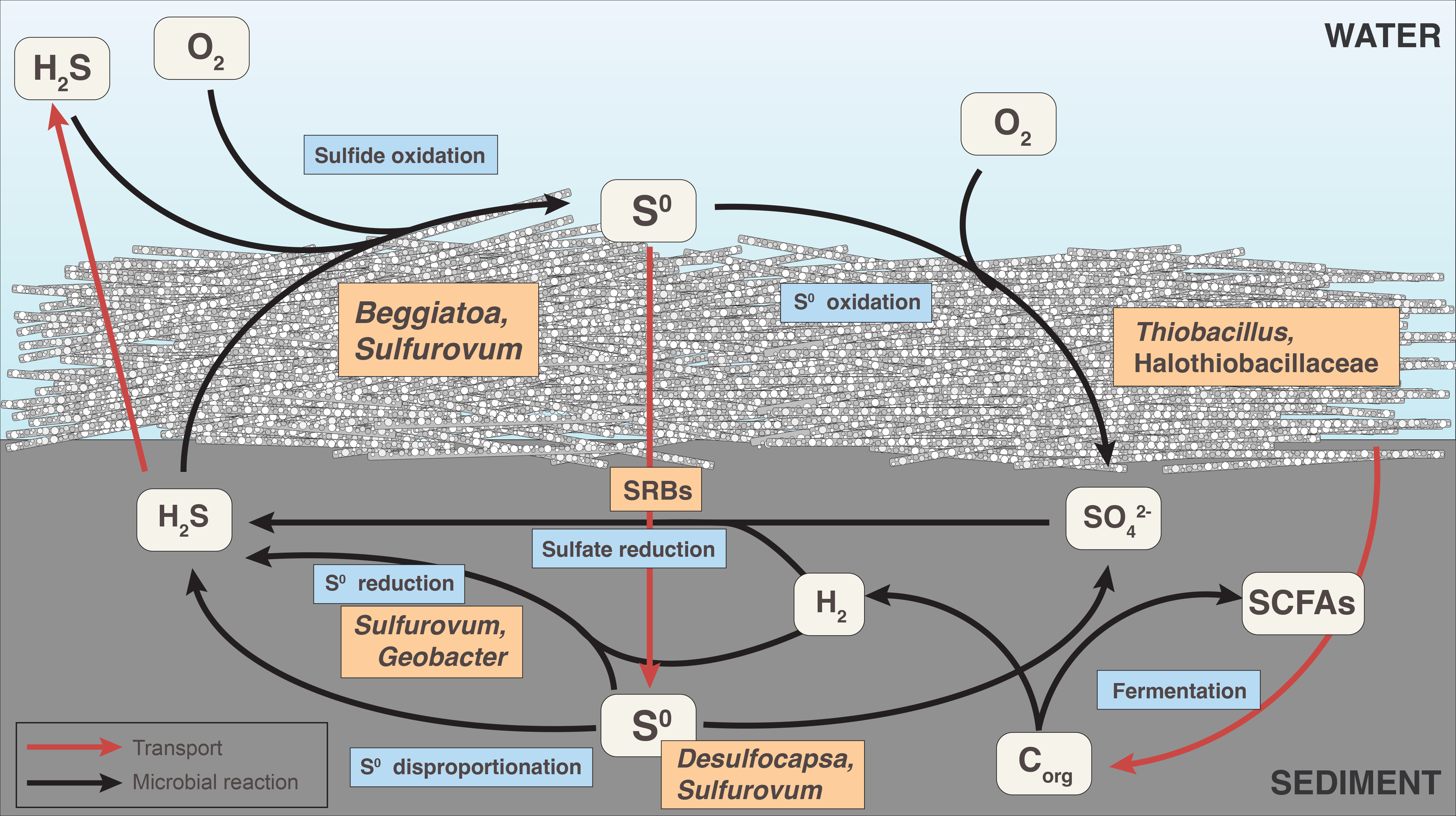
Suggested conceptual model of sulfur cycling in Frasassi biofilms and sediments. Blue boxes denote microbially catalyzed processes and orange boxes indicate the taxa responsible for them. In the *Beggiatoa* mats, incomplete sulfide oxidation to S^0^ is a dominant process, with possible minor contributions of sulfide or S^0^ oxidation with oxygen or nitrate. In the anoxic sediment, sulfide production may occur via S^0^ reduction or disproportionation, or sulfate reduction. SCFA stands for short chain fatty acids produced through fermentation reactions. The functions of microbes represented in the model were inferred from their closest cultured relatives and from previously-analyzed metagenome assembled genomes.

*Beggiatoa* spp., which dominated the mat community at 20-30% relative sequence abundances, have the genetic potential to oxidize reduced sulfur compounds (sulfide, S^0^, thiosulfate). *Beggiatoa* MAGs from Frasassi contain genes encoding enzymes involved in reduced sulfur oxidation, including *soxYZ, rDSR, fccAB,* and heterodisulfide reductases (McCauley Rench, 2015). While the presence of *rDSR* suggests that *Beggiatoa* spp. at Frasassi have the capacity to oxidize sulfide completely to sulfate (Mußmann *et al*., 2007), the availability of both sulfide and oxygen may determine the final product of sulfide oxidation (Klatt & Polerecky, 2015; Klatt *et al*., 2016). In marine and freshwater environments, *Beggiatoa* spp. have been shown to migrate vertically to take advantage of optimal geochemical parameters at oxygen and sulfide diffusion gradient zones. Under low sulfide flux conditions for a given O2 supply, *Beggiatoa* oxidize sulfide directly to sulfate, while under high sulfide flux conditions, *Beggiatoa* are only able to oxidize ∼50% of sulfide to sulfate, leading to intracellular S^0^ deposition (Berg *et al*., 2014) . Eventually, *Beggiatoa* may accumulate so much S^0^ that they burst (Berg *et al*., 2014). *Beggiatoa* may save themselves from this fate by migrating into anoxic/sulfidic regions where they can respire excess S^0^ using polyhydroxyalkanoates as electron donors (Schwedt *et al*., 2012). At Frasassi, *Beggiatoa* populations were not dependent on sulfide/oxygen supply ratios, but were observed in areas with less turbulent flow where they could accumulate on fine sediments (Macalady *et al*., 2008), perhaps indicating their ability to migrate to areas with ideal geochemical conditions. *Beggiatoa* MAGs from Frasassi also contained genes encoding cbb3-type cytochrome c oxidases and bd-type quinol oxidases, which are aerobic terminal oxidases that function under low oxygen conditions (Preisig *et al*., 1996; Rich *et al*., 1996; Pitcher & Watmough, 2004; Buschmann *et al*., 2010; McCauley Rench, 2015). In *E. coli*, cytochrome bd oxidases have been shown to promote sulfide-resistant respiration and growth and are expressed under microoxic conditions (Borisov *et al*., 2011, 2021; Forte *et al*., 2016). At Frasassi, *Beggiatoa* likely oxidize sulfide incompletely to S^0^ under the microoxic, sulfidic conditions of the streams, which is supported by the perennial observation of visible S^0^ inclusions within their cells and high concentration of dry weight sulfur in the cells (Fig. S1).

*Sulfurovum* spp., which were present at 5-15% relative sequence abundances across all samples, were most closely related to *Sulfurovum aggregans, Sulfurovum lithotrophicum, Sulfurovum riftiae* and *Nitratifractor salsuginis* (Fig. S5). These isolates are incapable of utilizing sulfide as an electron donor, and instead obligately oxidize H2 with S^0^, thiosulfate, or nitrate (*N. salsuginis* and *S. aggregans*; (Nakagawa *et al*., 2005; Mino *et al*., 2014)) or oxidize S^0^ or thiosulfate with oxygen or nitrate (*S. lithotrophicum* and *S. riftiae*; (Inagaki *et al*., 2004; Giovannelli *et al*., 2016)). However, prior analysis of four nearly complete *Sulfurovum* MAGs from Frasassi showed that these populations are capable of oxidizing sulfide and thiosulfate to S^0^ (*sqr* and *fcc*) using oxygen (cbb3-type cytochrome c oxidases and bd-type quinol oxidases) or nitrate (denitrification pathways) (Hamilton *et al*., 2015), suggesting that certain *Sulfurovum* spp. may contribute to S^0^ deposition within the microbial mats. It is also possible that different *Sulfurovum* populations in the sediment, which may be more closely related to cultured representatives, consume S^0^. Further cultivation and/or transcriptomic studies are necessary to determine the S^0^ consumption or production activity of *Sulfurovum* spp. in both the mats and sediments.

Uncultured strains related to the family Halothiobacillaceae were present at 2-10% across all samples and formed three clades with cultured relatives from the genera *Halothiobacillus*, *Thiovirga,* and *Thiofaba* (Fig. 3, Fig. S6)*. Halothiobacillus neapolitanus, Halothiobacillus kellyi, Thiovirga sulfuroxydans* and *Thiofaba tepidiphila*, the only cultured isolates from these genera, are autotrophs capable of complete aerobic sulfide, thiosulfate, and S^0^ oxidation to sulfate (Parker & Prisk, 1953; Sievert *et al*., 2000; Ito *et al*., 2005; Mori & Suzuki, 2008). Halothiobacillaceae from Frasassi mats have been shown to express the *sox* complex and *fccAB*, suggesting that these populations can oxidize sulfide and deposit S^0^, but also expressed *soxCD,* indicating that they completely oxidize intracellular S^0^ to sulfate (Pavia *et al*., 2018). In the mats, where oxygen is limiting and is rapidly scavenged to below detection limits (≤ 2 µM), some populations may oxidize sulfide or S^0^ using nitrate as an alternative electron acceptor. Such reactions yield -89 to -735 kJ/mol S (Table 1, Rxns 5-10), and could feasibly serve as alternative metabolisms when oxygen is limiting. Numerous taxa present in the mats, including *Beggiatoa* (Kamp *et al*., 2006; McCauley Rench, 2015)*, Sulfurovum* (Inagaki *et al*., 2004; Hamilton *et al*., 2015; Giovannelli *et al*., 2016; Pavia *et al*., 2018)*, Thiobacillus* (Schedel & Trüper, 1980), *Sulfuricurvum* (Kodama & Watanabe, 2004), and uncultured Halothiobacillaceae sequences closely related to ‘*Candidatus* Thiobacillus baregensis’ (Hédoin *et al*., 1996; Hédoin, 1997; Tsao, 2014) have close cultured relatives capable of sulfide, thiosulfate, and S^0^ oxidation with nitrate as an electron acceptor or have the genetic potential to perform this metabolism. The use of nitrate as an alternative electron acceptor could explain why nitrate levels at Frasassi are consistently below detection. However, the constant pH profile with depth through the microbial mat suggests that complete sulfide oxidation or S^0^ oxidation, either with oxygen or nitrate, is not a predominant sink for S^0^.

### S^0^ reduction and disproportionation in anoxic sediments

With increasing depth into the anoxic sediment (Fig. 1), a significant pH decrease (from 7.3 to 6.8) was observed and sulfide concentrations reached almost 600 µM at a depth of 3 mm below the mat-water interface (Jones *et al*., 2015)). Upward diffusion of sulfide from the anoxic sediments can explain the increasing sulfide concentration within the microbial mat (Fig. 1), contributing to the dynamic cycling of sulfide that occurs at Frasassi. Microbial metabolisms that generate sulfide include sulfate and S^0^ reduction or disproportionation of intermediate oxidation state sulfur species (sulfite, polythionates, thiosulfate, S^0^, or polysulfides) (Table 1, Rxns 11-16). Sulfur disproportionation, or the inorganic fermentation of sulfur, is the simultaneous oxidation and reduction of an intermediate oxidation state sulfur compound to form sulfate and sulfide. Under sulfidic conditions, polysulfides form through the reaction of S^0^ and sulfide, and concentrations of polysulfides increase proportionally with sulfide concentration (Schwarzenbach & Fischer, 1960; Schauder & Müller, 1993; Milucka *et al*., 2012). Under *in situ* conditions (pH ∼7, ∼500 µM sulfide), 47 µM of sulfide should be present as polysulfides, with Sn = 2, 4, 5, and 6 as the most abundant forms (Milucka *et al*., 2012), so it is possible that both polysulfides and S^0^ are disproportionated by microorganisms in Frasassi sediments.

Based on the microsensor data, sulfate reduction with H2 is more exergonic than S^0^ disproportionation or reduction (Δ*Gsulfate reduction* = -34 to -42 kJ/mol S, Fig. 2). However, when calculated per mole of H2, which is likely limiting in the sediments compared to S^0^, sulfate reduction yields only -8 to -10 kJ/mol H2 while sulfur reduction yields -19 to -29 kJ/mol H2 (Table 1). Although sulfate-reducing taxa including *Desulfonema, Desulfobacca,* and *Desulfatirhabdium* were detected in the amplicon dataset in low abundance (Fig. 3), sulfate reduction consumes rather than produces protons (Table 1, Reaction 5). Thus, while sulfate reduction was likely occurring in the sediments and could explain some of the increase in sulfide concentration, it was not occurring at rates high enough to counteract competing processes that produce protons. The decrease in pH with increasing depth in the anoxic sediments may also be partially attributed to short chain fatty acid production through fermentation by anaerobic chemoheterotrophs (Fig. 3, 4).

We therefore evaluated which sulfide-producing processes would also produce significant acidity. S^0^ reduction and disproportionation of S^0^ and polysulfides generate sulfide and protons, and in contrast to S^0^ oxidation, can proceed in the absence of oxidants and are therefore more likely to be sinks for S^0^ produced at the sediment-water interface. Taxa closely related to isolates that can reduce S^0^ in culture (*Sulfurovum* (Campbell *et al*., 2006; Mino *et al*., 2014)*, Sulfuricurvum* (Kodama & Watanabe, 2004; Campbell *et al*., 2006), uncultured Arcobacteraceae (Campbell *et al*., 2006), and *Geobacter* (Caccavo *et al*., 1994)) were abundant in the sediments. While S^0^ reduction with hydrogen yields more free energy per mole of sulfur than S^0^ disproportionation, S^0^ reducers are likely in competition for H2 with sulfate reducers. This could allow S^0^ and polysulfide disproportionating taxa to reach significant populations in the sediment because they are not in direct competition with sulfate reducing bacteria. Cultivated species of the genus *Desulfocapsa* are strictly anaerobic and couple autotrophic growth to the disproportionation of intermediate oxidation state sulfur species (S^0^, sulfite, and thiosulfate) to form sulfide and sulfate (Janssen *et al*., 1996; Finster *et al*., 1998, 2013; Frederiksen & Finster, 2003, 2004; Poser *et al*., 2013). Recently it was shown that several species of *Sulfurovum* from diverse natural environments carry out S^0^ disproportionation in pure culture (Wang *et al*., 2022). It is therefore possible that *Sulfurovum* populations are also carrying out S^0^ disproportionation at Frasassi, although their S^0^-disproportionating activity should be confirmed with laboratory studies. *Desulfocapsa* spp. and *Sulfurovum* spp. were found both in *Beggiatoa* mats and in sediments at 5-10% and 5-12% relative abundance, respectively, making up a total of 12-26% relative abundance in all samples (Fig. 3). Additionally, a novel family, genus, and species of S^0^-disproportionating bacteria was recently isolated from the Frasassi sediments (Aronson *et al*., 2022). Sequences associated with this strain were present at 0.11% relative abundance in the GS10-5-mat sample, suggesting that both rare and abundant members of the sediment and mat communities are potentially capable of S^0^ disproportionation. The presence of diverse sulfur disproportionating taxa and the geochemical trends in the sediments indicate that disproportionation is an important sink for S^0^ at Frasassi.

There are few studies that attribute polysulfide disproportionation to microbial processes (Milucka *et al*., 2012; Poser *et al*., 2013; Findlay, 2016). Based on our calculations, disproportionation of S2^2-^ is more exergonic than S^0^ disproportionation, S^0^ reduction, and sulfate reduction at Frasassi and could also be a significant contributor to sulfide and proton production in the sediments. Since S^0^ is a solid that can’t pass through cell membranes and polysulfides readily form by reaction of S^0^ and sulfide, it is likely that some S^0^ disproportionation occurs via a polysulfide intermediate produced either abiotically or biologically. While the biochemistry of S^0^ disproportionation is still poorly understood, environments with sufficient S^0^ and sulfide to produce abundant polysulfides may be especially important habitats for organisms carrying out disproportionation. *Sulfurovum* and *Sulfuricurvum* MAGs contained genes encoding polysulfide reductase (*psr*) and may be important contributors to polysulfide cycling in the Frasassi sediments (Hamilton *et al*., 2015).

In contrast to sulfur cycling in marine sediments, which is primarily driven by dissimilatory sulfate reduction to sulfide, S^0^ and polysulfide disproportionation are likely the predominant sulfur reactions in the anoxic Frasassi sediments. Despite numerous claims to the contrary (Bak & Cypionka, 1987; Thamdrup *et al*., 1993; Lovley & Phillips, 1994; Finster, 2008; Finster *et al*., 2013; Poser *et al*., 2013; Morrison & Mojzsis, 2021), S^0^ disproportionation is an energetically favorable process under a range of natural conditions (Alain *et al*., 2022). Erroneous claims of its endergonicity arise from confusion over what the term “standard state” means in thermodynamics (Amend & LaRowe, 2019). Thus, although the free energy yields of S^0^ disproportionation under *in situ* conditions are low, they clearly provide enough energy to sustain significant and diverse populations of S^0^ disproportionating microorganisms in the Frasassi sediments.

Certain laboratory cultures of disproportionators show increased growth in the presence of a sulfide scavenger such as Fe(III) (Thamdrup *et al*., 1993; Janssen *et al*., 1996; Finster *et al*., 1998). In these reactions, sulfide produced by sulfur disproportionation reduces FeOOH to form Fe^2+^ and S^0^ (Table 1, Rxn. 17). Sulfide reacts with Fe^2+^ to form FeS (Rxn. 18), giving Rxn. 19 as the overall reaction stoichiometry. Rusty-red iron crusts can be observed near Frasassi streams (Fig. S2), and both ferrous and ferric iron were measured in microbial mats and sediments (Fig. S3), indicating the potential for FeS products that could sequester sulfide and perhaps increase growth of sulfur disproportionating organisms in the sediment. However, the increase in sulfide concentration with depth in the sediment indicates that iron is limiting in the sediment, as the fast reaction of sulfide with FeOOH would prevent sulfide from accumulating in the sediment. Future studies are needed to investigate the dynamics of iron cycling in the Frasassi caves and their potential impact on sulfur cycling microbial communities.

In addition to serving as a sink for S^0^ at Frasassi, S^0^ or polysulfide disproportionation may also contribute to the weathering of carbonate rocks and sediments in sulfur-rich environments. Subaerial sulfuric acid speleogenesis via the abiotic and biological oxidation of sulfide in sulfidic karst has been studied intensively (Stern *et al*., 2002; Macalady *et al*., 2007; Jones *et al*., 2014). At Frasassi, subaqueous sulfuric acid speleogenesis is thought to have a minor contribution to cave formation because sulfide degassing rates exceed the rates of microbial sulfide oxidation, and because S^0^ rather than sulfuric acid is the major end product of subaqueous microbial sulfide oxidation (Jones *et al*., 2015). This is likely due to O2:H2S availability in the oxygen-rich cave air, where sulfide oxidation can proceed completely to sulfuric acid, while it is more limited by O2 availability in the microoxic streams. S^0^ disproportionation, which is an acid-producing process, may prove to be a significant contributor to anoxic, subaqueous sulfuric acid speleogenesis at Frasassi and in other sulfidic karst environments. This could be tested by incubating limestone tablets in the sediment, stream water, and cave air and measuring average mass loss over time (Peterson, 1966; Galdenzi, 2012; Jones *et al*., 2015), or by comparing rates of S^0^ disproportionation using radiolabeled compounds in field incubations.

### Frasassi sulfidic sediments as a Proterozoic analog environment

Microbial consumption of S^0^ has been studied in modern marine sediments (Pjevac *et al*., 2014). In sulfidic marine environments, including tidal flats and deep-sea hydrothermal vent systems, oxidation in surface sediments by *Sulfurimonas* and *Sulfurovum* was the main sink for S^0^. In anoxic sediments, *Desulfocapsa* colonized and consumed S^0^, indicating that disproportionation could be a sink for S^0^ in the absence of oxygen in marine sediments (Pjevac *et al*., 2014).

Sulfur cycling at Frasassi, which occurs under oxygen-limited conditions with significantly lower salinity and sulfate concentrations than modern oceans, may have significant similarities to Proterozoic sulfur cycling in fresh to brackish sediments, when euxinic marine ecosystems encountered the initial rise of oxygen (Meyer & Kump, 2008; Lyons *et al*., 2009). The fate of marine sedimentary S^0^ has likely varied in Earth history as the redox chemistry of the ocean evolved, and sulfur isotope fractionations preserved in the geologic record may reveal shifts in the predominance of competing sulfur metabolisms. In particular, the S^0^ disproportionation is thought to be an ancient catabolism based on phylogenetic and isotopic evidence (Canfield & Raiswell, 1999). Sulfur disproportionation produces sulfide depleted in ^34^S and sulfate enriched in ^34^S, and repeated cycles of sulfide oxidation to S^0^ followed by disproportionation produces sulfides that are strongly isotopically depleted (Canfield & Thamdrup, 1994; Canfield *et al*., 1998; Johnston *et al*., 2005). The closely linked ecology of sulfide oxidizers, sulfate reducers, S^0^ reducers, and S^0^ disproportionators at Frasassi may inform on how microbial metabolisms shaped the biogeochemical sulfur cycle in Proterozoic sedimentary ecosystems. At Frasassi, the *δ*^34^S fractionations observed between sedimentary sulfate and sulfides were smaller than what is expected in systems driven by reductive or disproportionative sulfur 16 cycling (Zerkle *et al*., 2016). Further studies are necessary to interpret the effects of chemolithotrophic sulfur oxidation on sulfur isotopic fractionation observed at Frasassi.

## Acknowledgements

The authors thank A. Montanari for providing logistical support and the use of facilities and laboratory space at the Osservatorio Geologico di Coldigioco, Apiro, Italy; S. Mariani, S. Cerioni, M. Mainiero, F. Baldoni, S. Carnevali and members of the Gruppo Speleologico C.A.I. di Fabriano for technical assistance during field campaigns, and to S. Dattagupta, R. McCauley Rench, KS Dawson and C. Chan for assistance in the field. We thank DS Jones for helpful discussions on Frasassi sediment biogeochemistry. This work was supported by National Science Foundation grants EAR-0525503 and EAR-1124411 (JM), the NASA Habitable Worlds program under grant 80NSSC20K0228 (DEL) and the USC Dornsife College of Letters, Arts and Sciences (DEL). HSA was supported by an NSF Graduate Research Fellowship grant DGE-1842487, the Lewis and Clark Fund for Exploration and Field Research in Astrobiology, the National Cave and Karst Institute NCKRI Scholar Fellowship, the Josephine de Karman Fellowship, and the Cave Conservancy Foundation Graduate Fellowship in Karst Studies.

## Data availability statement

16S rRNA amplicon reads are available in GenBank under BioProject accession number PRJNA871755. Chemical concentration data used for thermodynamic calculations are available in Supplemental File 1.

## Author contributions

HSA analyzed the data, interpreted results, designed the figures, and drafted the manuscript. CEC collected field samples, analyzed the data, interpreted results, designed the figures, and drafted the manuscript. DEL and JPA assisted with thermodynamic calculations and revised the manuscript. LP assisted with microsensor data collection and revised the manuscript. JLM designed the study and revised the manuscript. All authors approved the final version of the manuscript.

## Supplemental Figures

Figure S1: Total sulfur for mat and sediment samples from Frasassi cave streams (% dry wt.). Mean sulfur concentrations are 19% for biofilms and 2.3% for underlying sediments.

Figure S2: Photographs showing high (left) and low (right) stream flow at the Pozzo dei Cristalli sample location, Frasassi. There is no visible Fe(III) staining during high flow, but conspicuous Fe staining in sediments exposed to oxygen-rich cave air during low flow, as indicated by the red arrow.

Figure S3: Total Fe, Fe in silicates, pyrite Fe, Fe(III) in oxides, hydroxides and oxyhydroxides, and Fe(II) content of Frasassi stream biofilms and underlying sediments in % dry weight. Samples with sequential numbers are paired with each other (i.e. PC16-27b is a mat and PC16-28b is immediately underlying sediment). In the lower panel, triangles indicate sediment and circles represent *Beggiatoa spp.* dominated mats. X represents a streamer biofilm sample for comparison.

Figure S4: Values of Gibbs energies, Δ*Gr*, for elemental sulfur disproportionation as a function of sulfate and sulfide activity calculated according to 4S^0^ + 4H2O ⇌ 3HS^-^ + SO4^2-^ + 5H^+^ at 13.5°C and pH = 7.26. Sulfate activities were calculated using concentrations of 1x10^-6^ to 3x10^-3^ molal and sulfide activities were calculated using concentrations of 1x10^-6^ to 1x10^-3^ molal. The red ellipse represents the energetic yield per reaction of sulfur disproportionation under Frasassi sediment conditions (sulfate activity of 2x10^-3^ and sulfide activities ranging from 2x10^-6^ to 6x10^-4^).

Figure S5: Phylogenetic tree of 16S rRNA gene sequences affiliated with the genus *Sulfurovum* from this study, the NCBI database, and previous Frasassi studies (n = 107). Sequence names in bold are isolates. Numbers in parentheses indicate the average relative abundance of sequences in microbial mats and sediments, respectively. Percentages listed next to collapsed clades indicate the total relative abundance of sequences within the clade. Collapsed clades without numbers indicate clades composed entirely of sequences from previous Frasassi studies.

Figure S6: Phylogenetic tree of 16S rRNA gene sequences affiliated with the family Halothiobacillaceae and the genus *Thiobacillus* from this study, the NCBI database, and previous Frasassi studies (n = 94). Sequence names in bold are isolates. *Candidatus* Thiobacillus baregensis is located within the Frasassi *Thiofaba* collapsed clade. Numbers in parentheses indicate the average relative abundance of sequences in microbial mats and sediments, respectively. Percentages listed next to collapsed clades indicate the total relative abundance of sequences within the clade. The genus *Halothiobacillus* was originally part of the genus *Thiobacillus*, which contained species that were classified as *α*-, *β*-, and *γ*-Proteobacteria (Kelly & Wood, 2000). The genus *Halothiobacillus* has recently undergone reclassification, but originally contained four species (*H. neapolitanus, H. kellyi, H. hydrothermalis,* and *H. halophilus*) of obligately autotrophic, aerobic sulfide, thiosulfate, and S^0^ oxidizing bacteria (Boden, 2017). *H. hydrogthermalis* and *H. halophilus* have been reclassified to the genus *Guyparkeria* within the family Thioalkalibacteraceae, which also contains *Thioalkalibacter halophilus* (Banciu *et al*., 2008; Boden, 2017).

Figure S7: Phylogenetic tree of 16S rRNA gene sequences affiliated with the genus *Desulfocapsa* from this study, the NCBI database, and previous Frasassi studies (n = 43). Sequence names in bold are isolates. Numbers in parentheses indicate the average relative abundance of sequences in microbial mats and sediments, respectively. Percentages listed next to collapsed clades indicate the total relative abundance of sequences within the clade.

**Supplemental File 1:** Concentrations of aqueous chemical species from Pozzo dei Cristalli (Tab 1, “DepthProfile”). Concentrations were sourced from median *in situ* values collected between 2002 and 2011. Concentrations of sulfide, oxygen, and protons were sourced from microsensor data from Jones *et al*. 2015. Oxygen concentrations below the detection limit (≤2 µM) were set between 10^-6^ molal for depths 0.5 mm and 0.55 mm and 10^-8^ molal for depths below 0.6 mm. Concentration gradients were created for sulfate (Tab 2, “SulfateGradient”) and sulfide (Tab 3, “SulfideGradient”), with the concentrations of all other aqueous species remaining constant.

## Notes

### Competing Interest Statement

The authors have declared no competing interest.

