## Supplementary figures and images for "Sulfur disproportionating microbial communities in a dynamic, microoxic-sulfidic karst system"

### Supplemental Figures

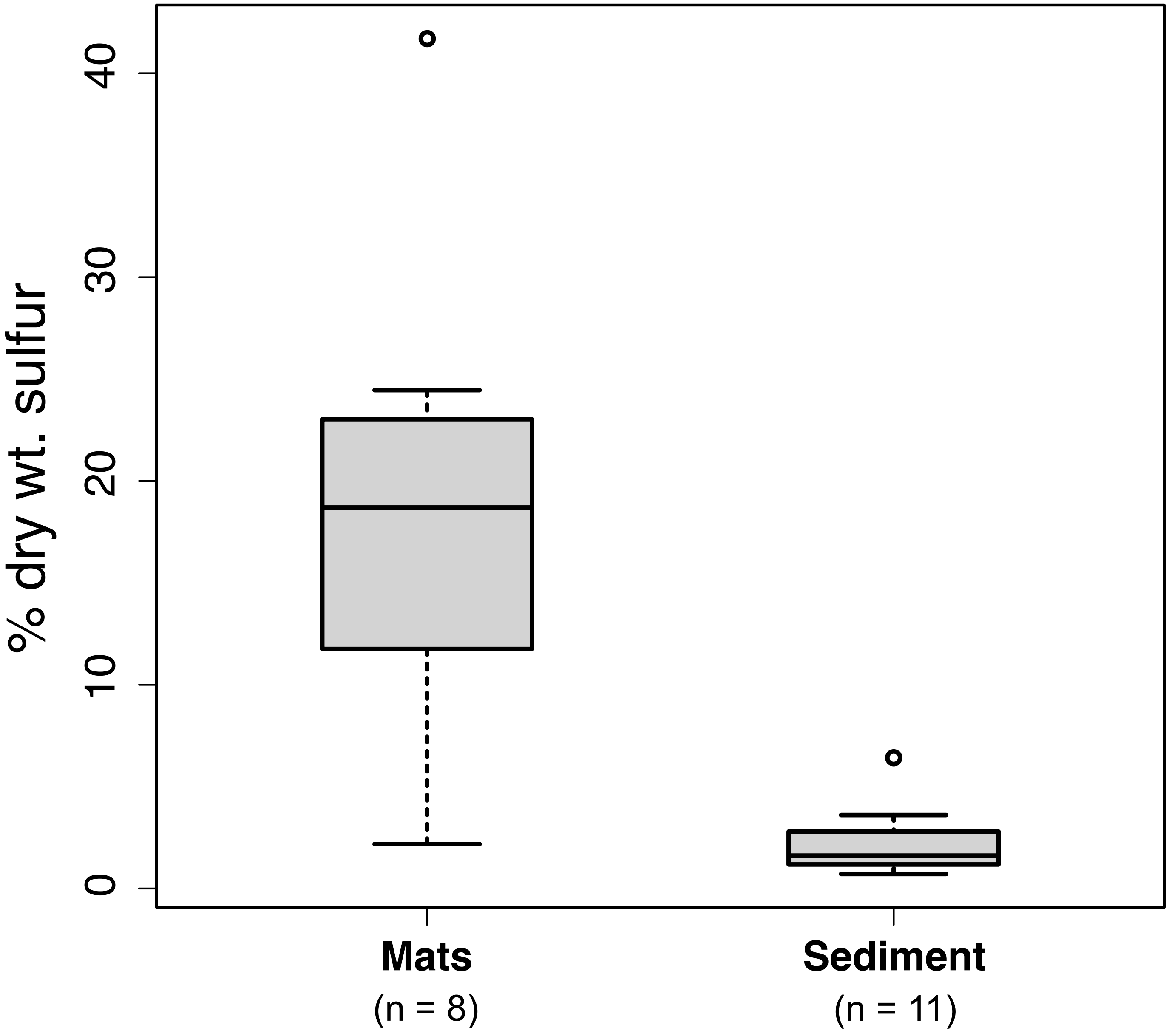

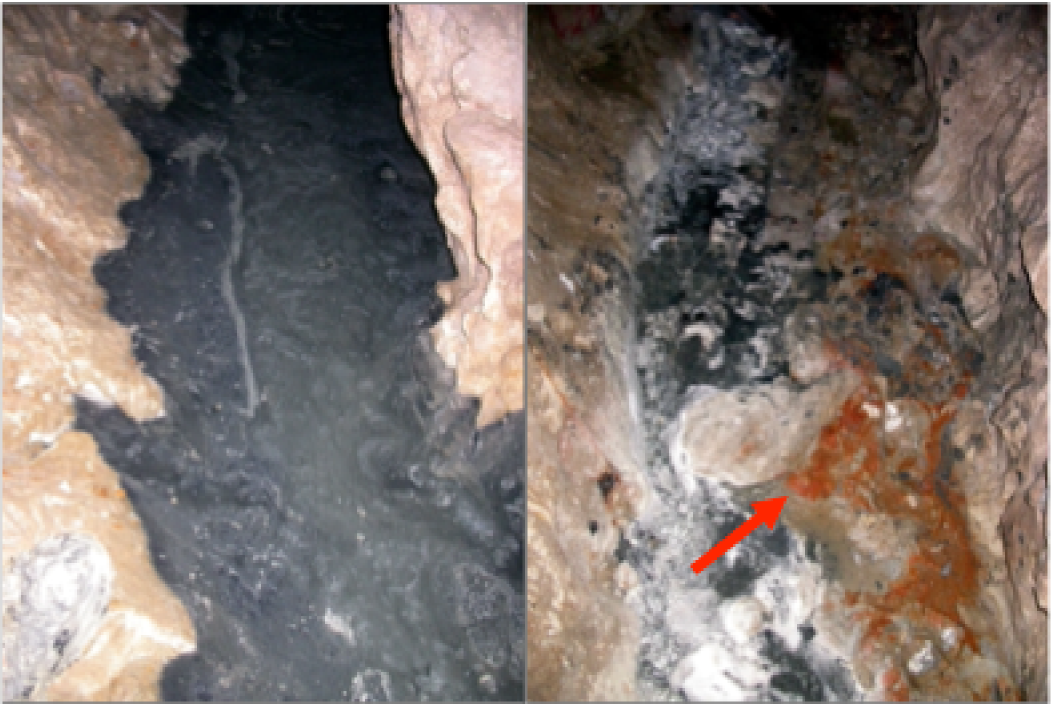

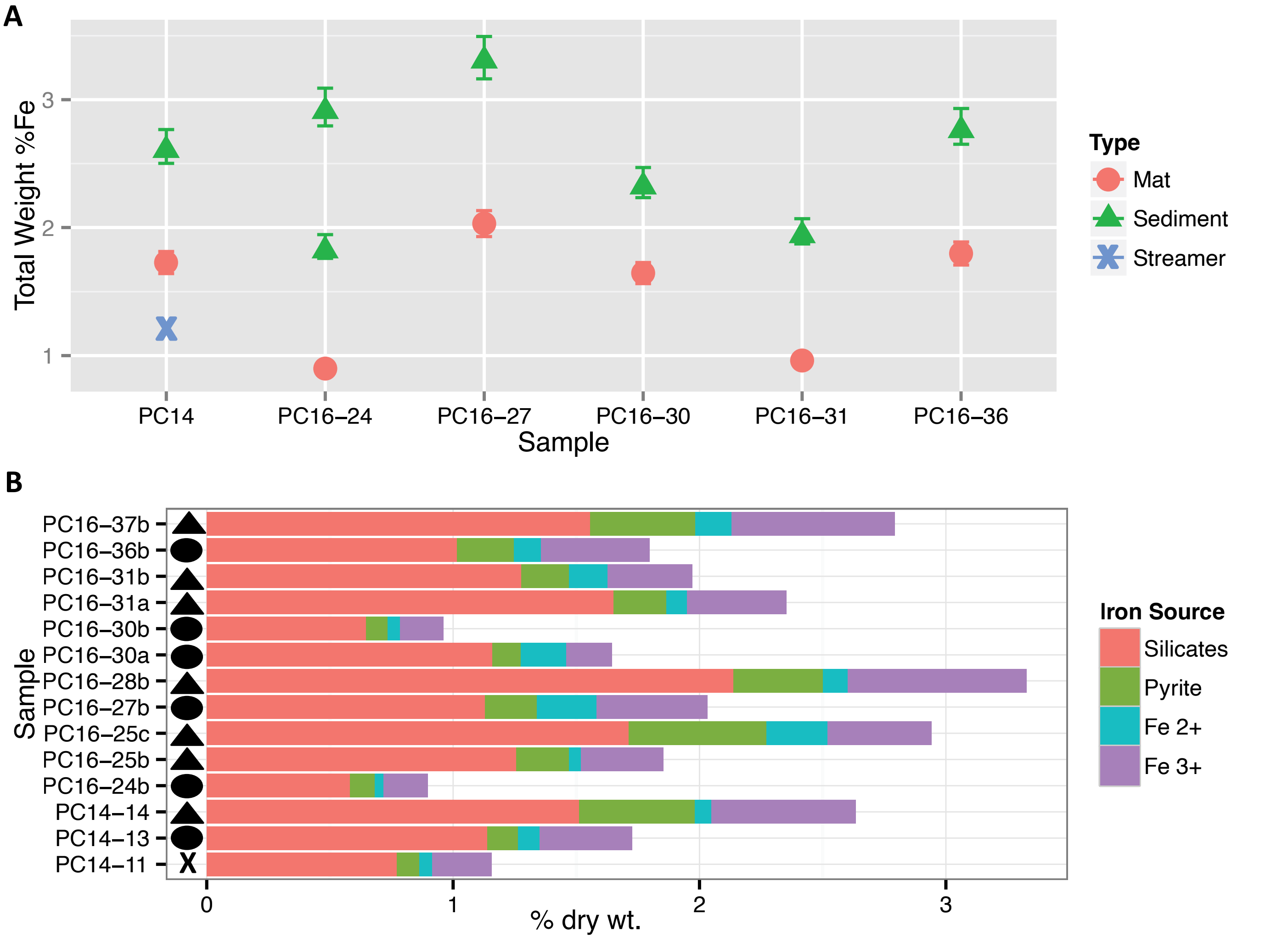

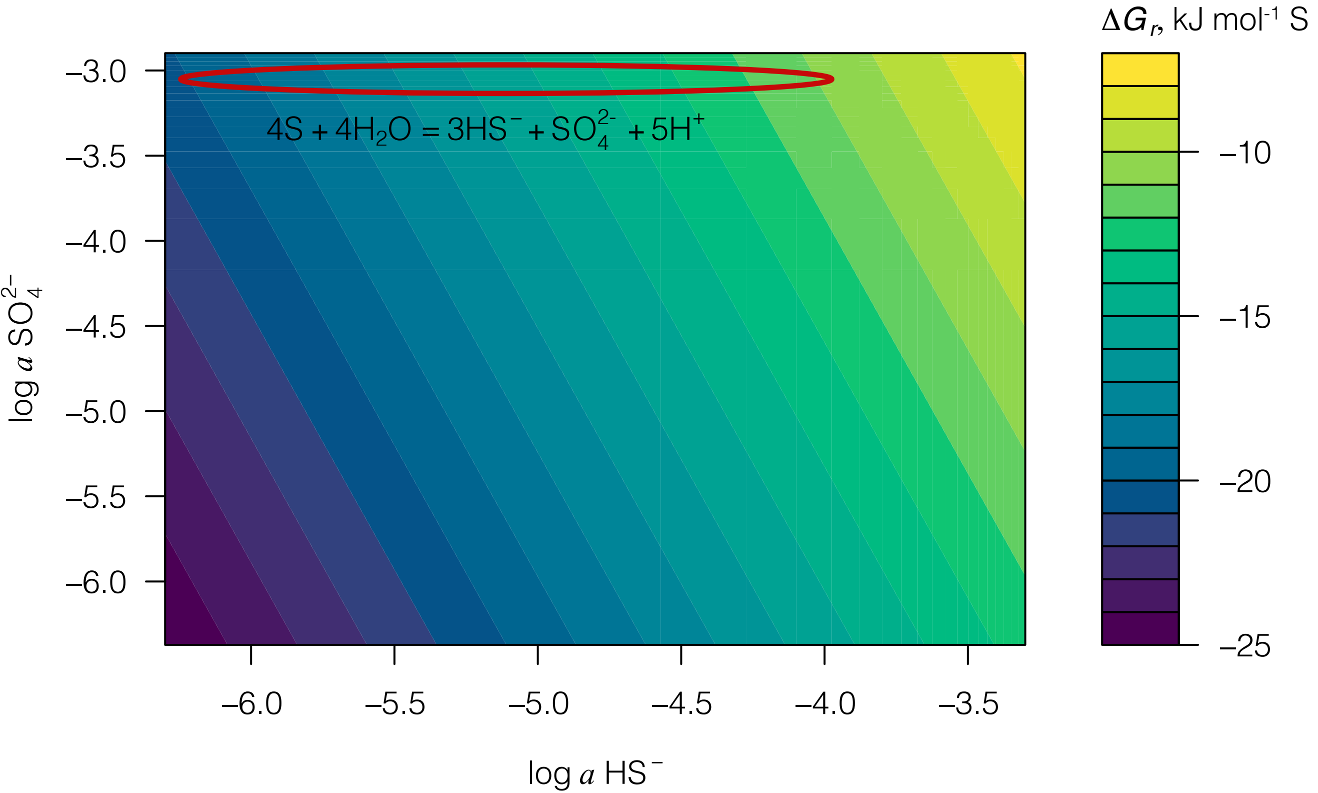

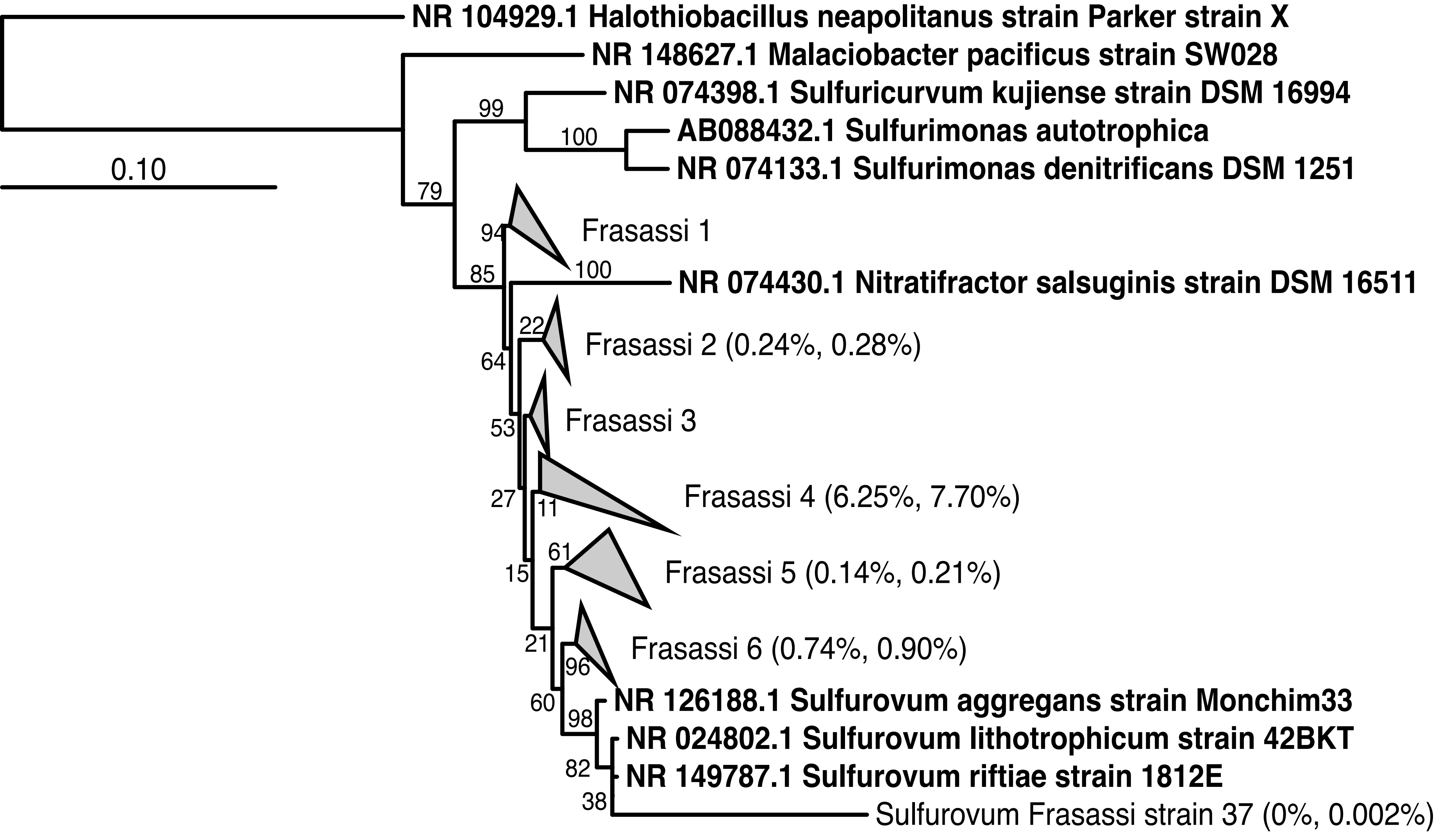

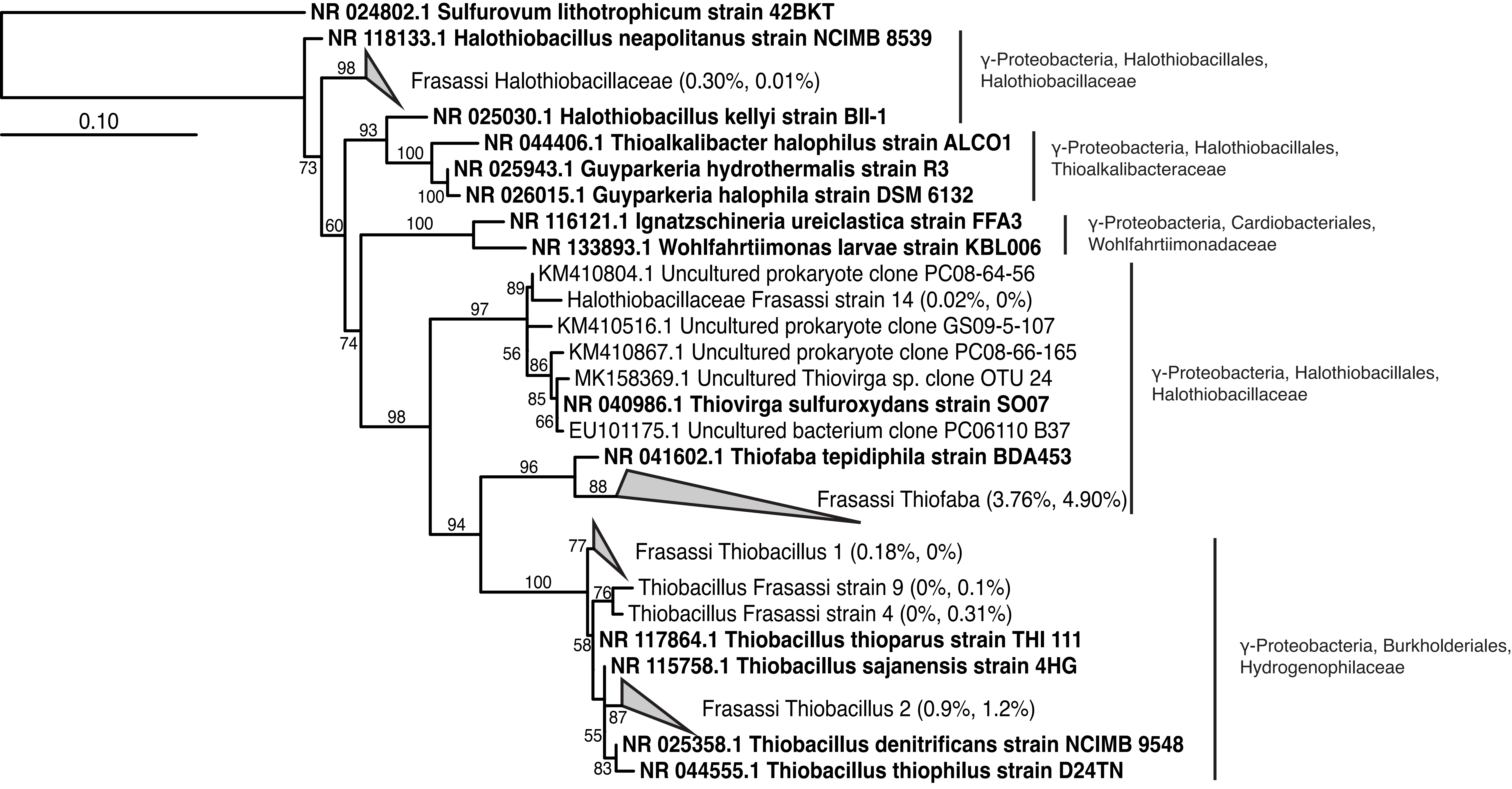

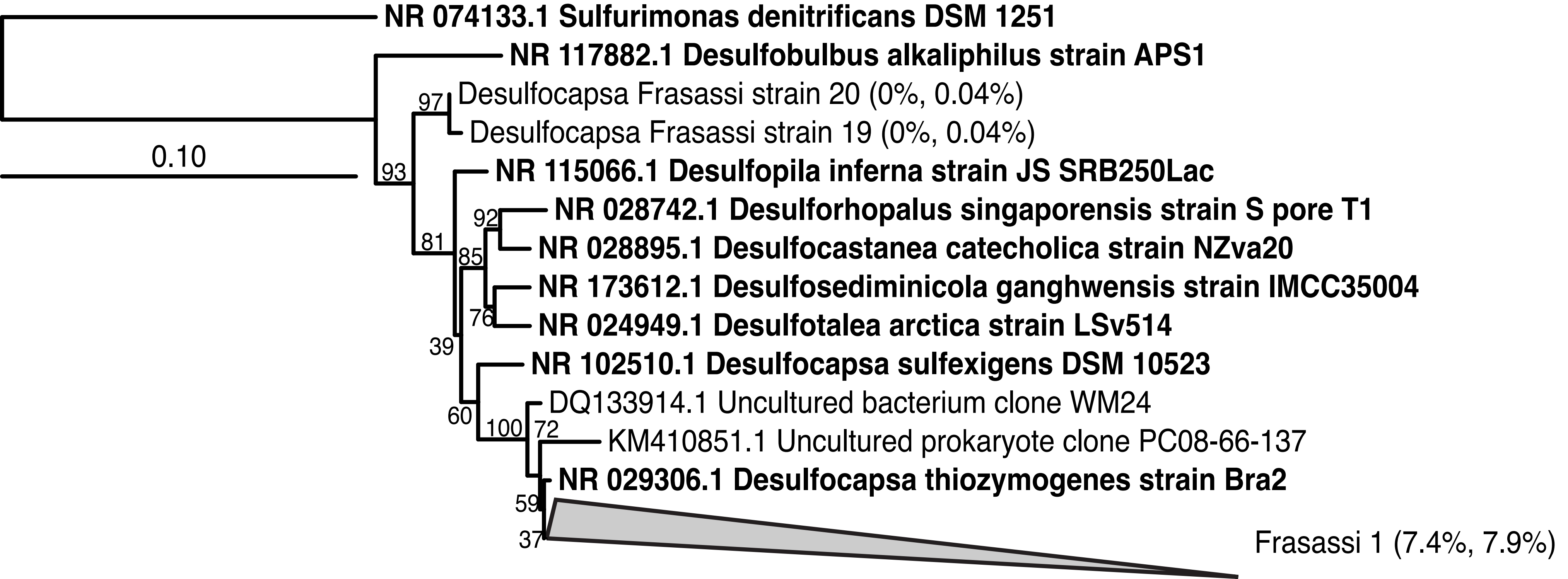
